# Modulation of ADARs mRNA expression in congenital heart defect patients

**DOI:** 10.1101/365288

**Authors:** Faiza Altaf, Cornelia Vesely, Abdul Malik Sheikh, Rubab Munir, Syed Tahir Abbass Shah, Aamira Tariq

## Abstract

Adenosine (A) to inosine (I) RNA editing, is a hydrolytic deamination reaction catalyzed by adenosine deaminase (ADAR) acting on RNA enzymes. RNA editing is a molecular process that involves the post-transcriptional modification of RNA transcripts. Interestingly, few studies have been carried out to determine the role of RNA editing in vascular disease. The current study found that in blood samples positive for congenital heart disease (CHD) ADAR1 and ADAR2 expression change at RNA level was opposite to each other. That is, an increase of ADAR1 mRNA was noticed in human CHD cases, whereas ADAR2 mRNA was vastly down-regulated. The increase in ADAR1 may be explained by the stress induced by CHD. The dramatic decrease in ADAR2 in CHD cases was unexpected and prompted further investigation into its effects on the heart. Therefore we performed expression analysis on a microarray data encompassing ischemic and non-Ischemic cardiomyopathy patient myocardial tissues. A strong down-regulation of ADAR2 was observed in both ischemic and especially non-ischemic cases. However, ADAR1 showed a mild increase in the case of non-ischemic myocardial tissues. To further explore the role of ADAR2 with respect to heart physiology. We selected a protein coding gene filamin B (FLNB). FLNB is known to play an important role in heart development. Although there were no observable changes in its expression, the editing levels of FLNB dropped dramatically in ADAR2^-/-^ mice. We also performed miRNA profiling from ADAR2 ^-/-^ mice heart tissue revealed a decrease in expression of miRNAs. It is established that aberrant expression of these miRNAs is often associated with cardiac defects. This study proposes that sufficient amounts of ADAR2 might play a vital role in preventing cardiovascular defects.

## Introduction

Congenital Heart Disease (CHD) is defined as structural or functional heart defect. It belongs to a heterogeneous group of diseases and can be classified anatomically, clinically, epidemiologically and developmentally (1-3). The most common types of CHDs among hospitalized patients are VSD (Ventricular Septal Defects), TOF (Teratology of Fallot), PDA (Patent Ductus Arteriosus), TGA (Transposition of Arteries), ASD (Atrial Septal Defect) and CAVSD (Atrioventricular Septal Defect) (4). Recent studies indicate that 11% of Pakistani children die due to cardiac anomalies at the first postnatal month (5).. Genetic conditions or environmental factors such as maternal diabetes or rubella are identified in some cases but for most babies born with a heart defect the cause remain unknown(6).

The multi-lineage differentiation during cardiogenesis is orchestrated by a precise spatial and temporal regulation of gene expression. Genetic studies in humans and knockout embryos have identified various genes, such as *TBX5*, *NKX2*-*5*, *GATA4*, *CX43*, *NOTCH1* and *VEGF* responsible for sporadic and inherited CHD cases (7).

In humans, the most prevalent type of RNA editing is adenosine (A) to inosine (I) (8). This complex post-transcriptional hydrolytic deamination reaction is carried out by adenosine deaminase (ADAR) family of enzymes. This family acts on double stranded RNA and comprise of three members ADAR1, ADAR2 and ADAR3. ADAR1 ad ADAR2 are actively involved in adenosine deamination however, ADAR3 is non-functional (9). Different studies have shown that the extent of RNA editing not only varies among individuals but also show high tissue specificity. Approximately 2.5 million sites in human transcriptome undergo editing however a vast majority of them lie in the Alu elements located mostly in the introns and UTR (untranslated region) (10). However, the functional consequence of majority of RNA editing events still remain elusive. RNA editing is known to modulate splicing, coding potential, transcript stability and even alters the processing and targeting of the microRNAs (miRNA) (8, 11, 12). RNA editing process affects RNA stability by conversion of a stable A:U base pair to a relatively unstable I:U base pair followed by unwinding of the RNA duplex and making it susceptible to single strand specific RNases (11). Recent study focusing on RNA editing events in six different tissues have demonstrated an average of 79,976 editing sites in heart (10). Previous report focusing on cyanotic congenital heart disease has indicated a significant decline in ADAR2 RNA level but no prominent difference in ADAR1 expression. Moreover, they showed high editing of MED13 in cyanotic CHD cases as opposed to acyanotic CHD patients (13). Recent study has shown an increase in expression of a lysosomal cysteine protease encoded by cathepsin S RNA (CTSS) via ADAR1 mediated RNA editing followed by HuR recruitment. Cathepsin S has a role in vascular inflammatory processes and the CTSS mRNA editing is increased in hypoxic or pro-inflammatory conditions as wells as in patients suffering from clinical or subclinical vascular damage (14).

In the current study we have determined the RNA level of ADAR1 and ADAR2 in congenital heart disease patients. We also checked the relative gene expression of FoxP1 which is an important transcription factor crucial for angiogenesis. We found a strong down-regulation of ADAR2 and an up-regulation of ADAR1. Interestingly, microarray data analysis of human non-ischemic myocardial tissues showed similar trend. Interestingly the ischemic myocardial tissues showed completely opposite trend. To further explore the role of ADAR2 in heart physiology, we used ADAR2 knockout mouse. Although no strong anomaly in heart physiology was observed as documented previously (15) however, we found down-regulation of different microRNAs.

## Materials and Methods

### Collection of samples

The blood samples were collected from 35 patients displaying different congenital heart defects from RIC (Rawalpindi institute of cardiology), stored in ice during transportation. Samples were segregated on the basis of age (3 months-16 years) and sex. Patient’s echocardiography reports were consulted to confirm the presence of congenital heart defects and all sample collection was done pre-operatively. Whereas 13 control samples were collected from healthy individuals using same parameters. Interviews were conducted personally using the specified questionnaires. Information on age, gender, medication and family history was recorded. Perspective study was initiated after getting approval from ethical committees of both CIIT and collaborating hospitals.

### cDNA synthesis

Five cubic centimeters of whole blood was collected from each patient in ethylenediamine tetraacetic acid (EDTA) test-tubes. To avoid RNA degradation, blood was kept at 4°C up to 24 h following collection before RNA was extracted. RNA extraction was carried out from peripheral blood mono-nuclear cells using the TRIzol ^®^ LS Reagent (Invitrogen, Germany) according to manufacturer’s instruction. Optical density of the RNA was measured immediately following extraction. RNA samples showing A260/280 below 1.8 or above 2.0 were not taken for further analysis. One microgram of RNA was used for production of complementary DNA (cDNA) using Revert aid first strand cDNA synthesis kit (Thermo scientific, USA). A negative control was set up against each of the sample that lacked Reverse transcriptase and was termed as –RT (minus Reverse Transcriptase).

### Real time PCR

The relative mRNA expression of genes were examined using a quantitative PCR with gene specific primer sets (IDT,USA and Macrogen, South Korea) and *TUBB11* was taken as internal control. 5x HOT FIREPol^®^ EvaGreen^®^qPCR Mix plus (ROX) (Solis Bio Dyne, Tartu, Estonia) master mixed was used for qPCR reaction. Sequence was taken from ensemble and primers were synthesized by Integrated DNA Technology (**biotools.idtdna.com/Primer Quest).** The primers are listed in the supplementary table 1

### Statistical analysis

Statistical analyses were performed with Graph-pad Prism 7.0b. For expression data, the target genes (*ADAR1*, *ADAR2*, *FOXP1*) CT was normalized with the control gene (*TUB1*) Ct. Depending on experiment, the statistical significance was determined using the Mann-Whitney test) with P< .05 considered significant.

### MicroArray analysis

Micro array data analysis was performed using CARMA web tool (16) on the GEO data set GDS1362 (17)focusing on expression analysis from myocardial tissues of non-ischemic (NICM), Ischemic cardiomyopathy (ICM) patients as opposed to non-failing heart tissues. The raw micro array data was extracted and normalization of the data was performed by gcRMA package.

### Mice

The *Adar2*^−/−^ knockout mouse was a kind gift of Peter Seeburg. These transgenic mice are in an SV129 background. As ADAR2 deficiency leads to early postnatal lethality, the mice were rescued with a pre-edited Gria2 receptor (*Gria2^R/R^*) (18, 19). Mice were bred in Vienna Biocentre facility animal house. *Gria2^R/R^; ADAR2*^+/−^ were intercrossed. The resulting sibling female offspring of genotype *Gria2^R/R^; Adar2*^−/−^ and *Gria2^R/R^; ADAR2*^+/+^ was euthanized at the age of post natal day 6 (P6). Whole heart was dissected and subsequently used for RNA preparation from three biological replicates (18, 20).

### RNA extraction and miRNA cloning

Female mouse whole heart as dissected at the age of post natal day 6 (P6), homogenized and total RNA was extracted using TriFast reagent according to manufacturer’s instructions (PEQLAB Biotechnologie GmbH, Erlangen, Germany). miRNA library preparation was performed as previously described (21).

### Sequencing and clipping of reads

Completed libraries were quantified with the Agilent Bioanalyzer dsDNA 1000 assay kit and Agilent QPCR NGS library quantification kit. Cluster generation and sequencing was carried out using the Illumina Genome Analyzer IIx system according to the manufacturer’s guidelines. Illumina sequencing was performed at the CSF NGS Unit (csf.ac.at). After sequencing at a read length of 36 base pairs, adaptor sequences were removed using Cutadapt (22).

### Mapping to mature miRNA sequences

Mapping of clipped reads to mature miRNA sequences was performed as described previously Mapping was performed using NextGenMap, restricting the mapped reads to have at least 90% identity (# differences/alignment length) (23).

## Results

### ADAR2 has the lowest expression in CHD patients

RNA was extracted from 35 affected congenital heart disease patients and 13 normal individuals were taken as control. Most of the patients had VSD (Ventricular septal defects). All CHD samples were pre-operative cases. Since recent reports indicated, that RNA editing might be involved in cardiogenisis (13, 14). We checked the expression of the functional RNA editing enzymes using quantitative realtime PCR (qPCR). A significant decline in ADAR2 expression was observed. However, on the contrary, ADAR1 showed a significant increase (Figure 1a). The observed elevated ADAR1 expression is in line with the recent finding demonstrating higher ADAR1 expression in the human patients undergoing carotid endarterectomy operation(14). As a control for CHD, we selected a member of fork head box family of transcription factor (FoxP1). FoxP1 has a critical role in murine as well as human heart development. It showed high expression during embryonic stages as opposed to postnatal stage and is critical for cardiomyocyte proliferation (24). We found a significant decrease in FoxP1 expression confirming that the RNA extracted from the blood of CHD patients can depict the expression differences between normal and patient samples. Surprisingly out of these three genes, ADAR2 was strongly down-regulated (Figure 1) pointing towards its expression modulation in relation to heart disease.

**Figure 1.**
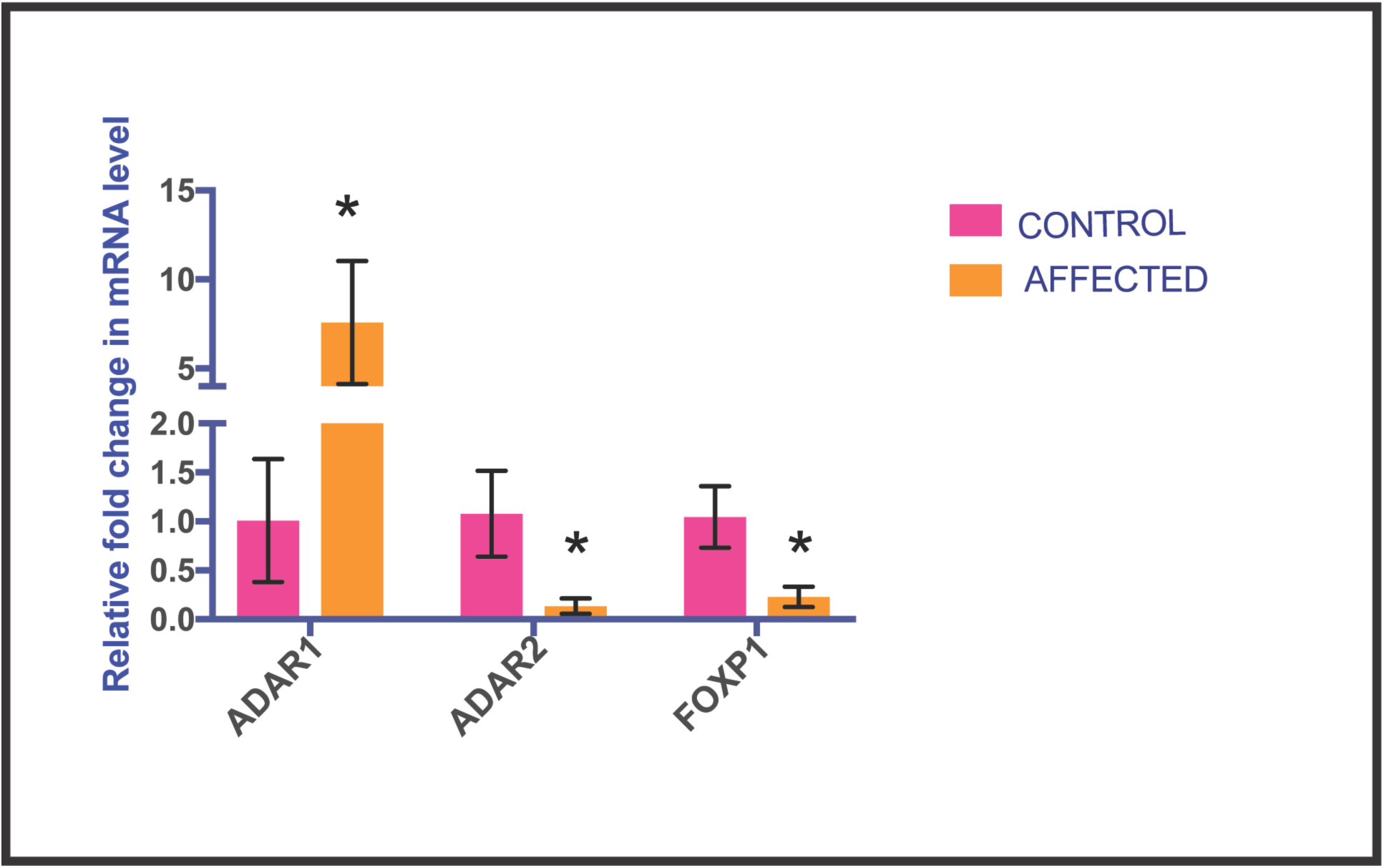
ADAR2 has the lowest expression in CHD patients: Bar graph showing relative RNA level of ADAR1, ADAR2 and FOXP1 as opposed to control. ADAR2 and FOXP1 show a significant reduced expression at RNA level whereas ADAR1 shows upregulation. ^∗^*P* < 0.05 (versus control) was determined by Mann–Whitney *U* test.

### Heart defect specific function

We further investigated whether this increase in ADAR1 is specific for a heart defect. As in our study the patients were suffering from different forms of congenital heart disease. ADAR1 was strongly up-regulated in ASD followed by VSD. However, it shows approximately 3fold increase-in TOF and CAVSD (Figure 2a and d). On the contrary, ADAR2 shows a strong significant decline in CAVSD, TOF and VSD. (Figure 2b and d). This specificity of gene expression of ADAR1 and ADAR2 with heart defects can be answered by the differences in the cardiac myocytes. Atrial, ventricular and nodal cells are morphologically, molecularly and functionally distinct. We propose that ADAR1 might have a more critical function in atrium development as compared to ventricles whereas ADAR2 might be more crucial for ventricular development. In the case of FOXP1, the most prominent decline was observed in the TOF patient samples (Figure 2c and d). FOXP1 plays a critical role in maintaining a balance of cardiomyocyte proliferation and differentiation via regulation of Fgf ligand and modulation of Nkx2.5 expression (25).This might answer the observed an increase in ADAR1 mRNA level and a strong decrease in FOXP1 expression.

**Figure 2.**
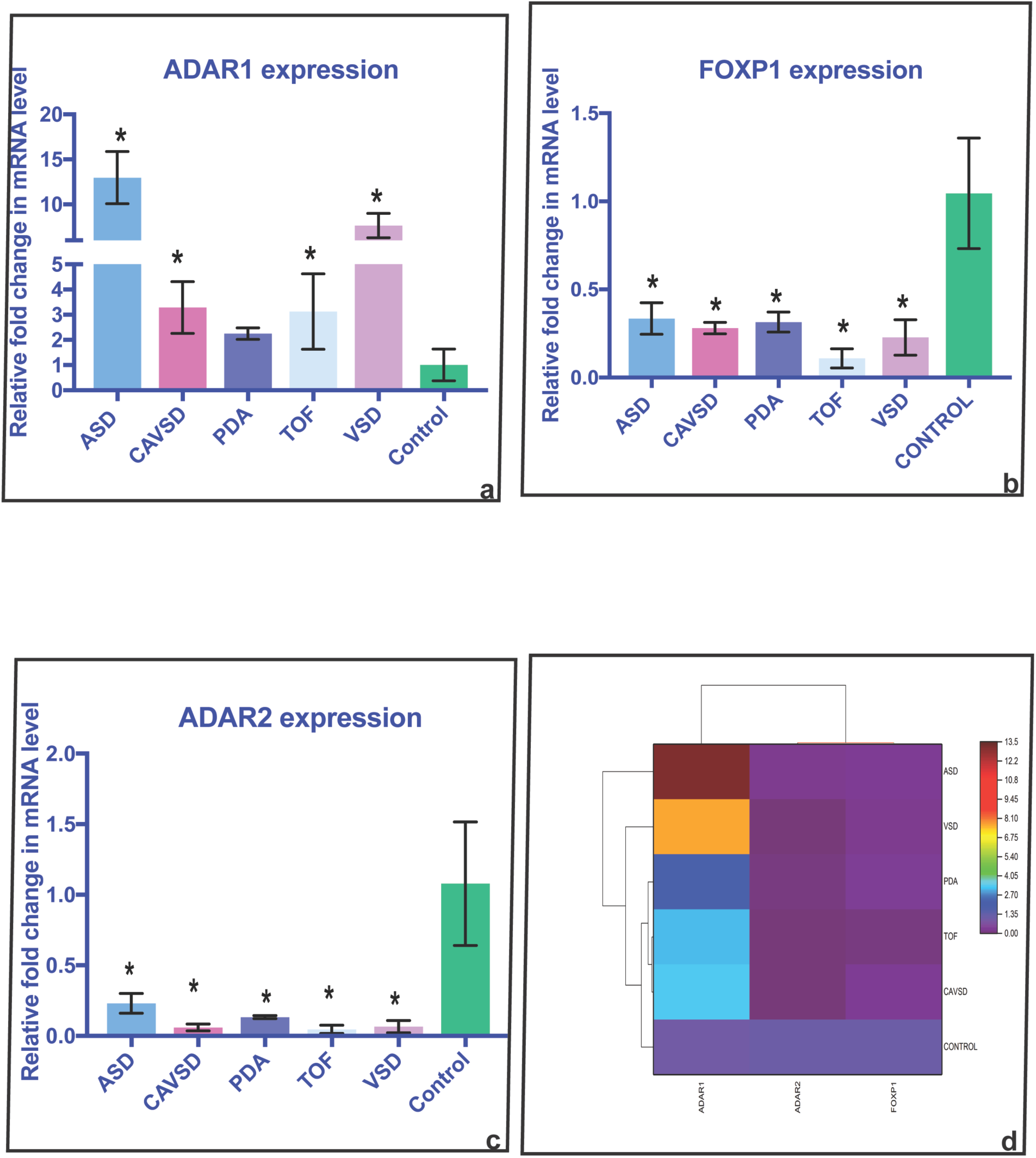
Heart defect specific function. a. Bar graph showing up-regulation of ADAR1 in different CHD cases. ADAR1 is significantly up-regulated in ASD and VSD. However, 3-4 fold increase at RNA level was found in AVSD and TOF cases. The PDA cases did not show a significant change. ^∗^*P* < 0.05 (versus control) was determined by Mann–Whitney *U* test.
b. Bar graph showing strong decline in ADAR2 expression particularly in AVSD, TOF and VSD. ADAR2 is significantly down-regulated in all CHD cases. ^∗^*P* < 0.05 (versus control) was determined by Mann–Whitney *U* test.
c. Bar graph showing reduced expression of FOXP1 in CHD cases. The strongest decline was observed in TOF samples. ^∗^*P* < 0.05 (versus control) was determined by Mann–Whitney *U* test.
d. Heat map showing expression level of ADAR1, ADAR2 and FOXP1 as opposed to control in different CHD cases.

### ADAR1, ADAR2 and Cardiomyopathy

Since all the above-mentioned results, were observed only in PBMCs we performed microarray data analysis on the Geo Dataset GDS1362 (17). This dataset comprised of differentially expressed genes in myocardial tissues from non-failing heart, ischemic and non-ischemic cardiomyopathy patients. We found a similar trend of expression of ADAR1 and ADAR2 in myocardial tissues from non-ischemic patients. ADAR1 showed a slight but significant increase whereas ADAR2 showed a strong decline in expression. Interestingly we found that the decrease in expression of ADAR2 was more prominent in non-ischemic as opposed to ischemic myocardial tissues (Figure 3).

**Figure 3.**
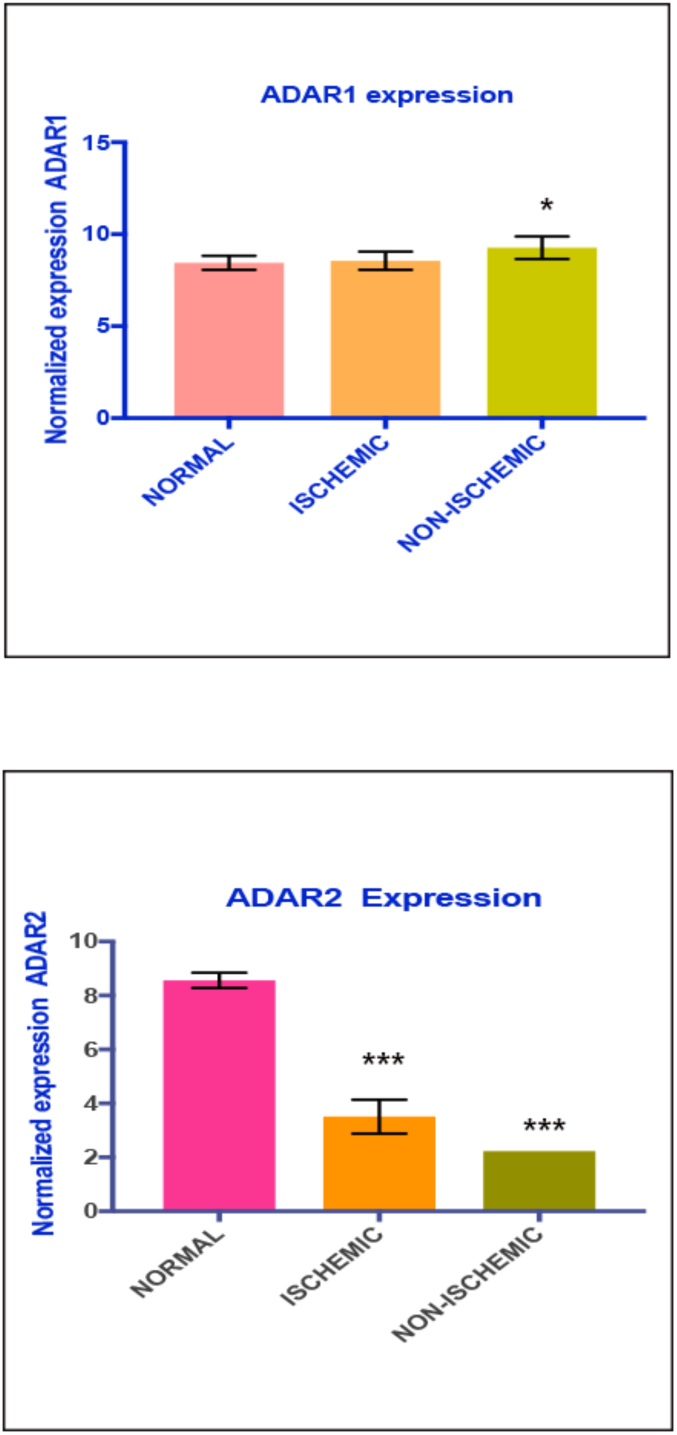
**(a)** Bar graph showing slight increase in ADAR1 expression in ischemic myocardial tissues from cardiomyopathy patients. **(a)** Bar graph showing significant decrease in ADAR2 expression in ischemic and non-ischemic myocardial tissues from cardiomyopathy patients. ^∗^*P* < 0.05 (versus control) was determined by Mann–Whitney *U*test.

### Filamins and Cardiac Defects

The strong decrease of ADAR2 expression in CHD patients made us curious to further investigate what happens in ADAR2^-/-^ mice heart?. Actin binding proteins such as FLNA and FLNB play essential role in the vascular development. FLNA is ubiquitously expressed whereas FLNB expression is mainly in the endothelial cells (26). Complete loss of FLNA results in severe structural defects in the heart involving atria, ventricles and outflow tracts (27). The decrease in ADAR2 mRNA level in CHD patients has urged us to look for protein coding targets that play significant role in cardiac development. Therefore, we chose FLNB. FLNA has been previously reported as ADAR2 editing target in the heart. Editing of FLNA drops down dramatically in ADAR2^-/-^mice heart (28). FLNB shows highest editing in the heart as compared to other tissues (28). Therefore, we checked the expression and editing of FLNB in the absence of ADAR2. We observed a dramatic decrease of 24% in editing of FLNB in ADAR2^-/-^ mice heart (Figure 4).

**Figure 4.**
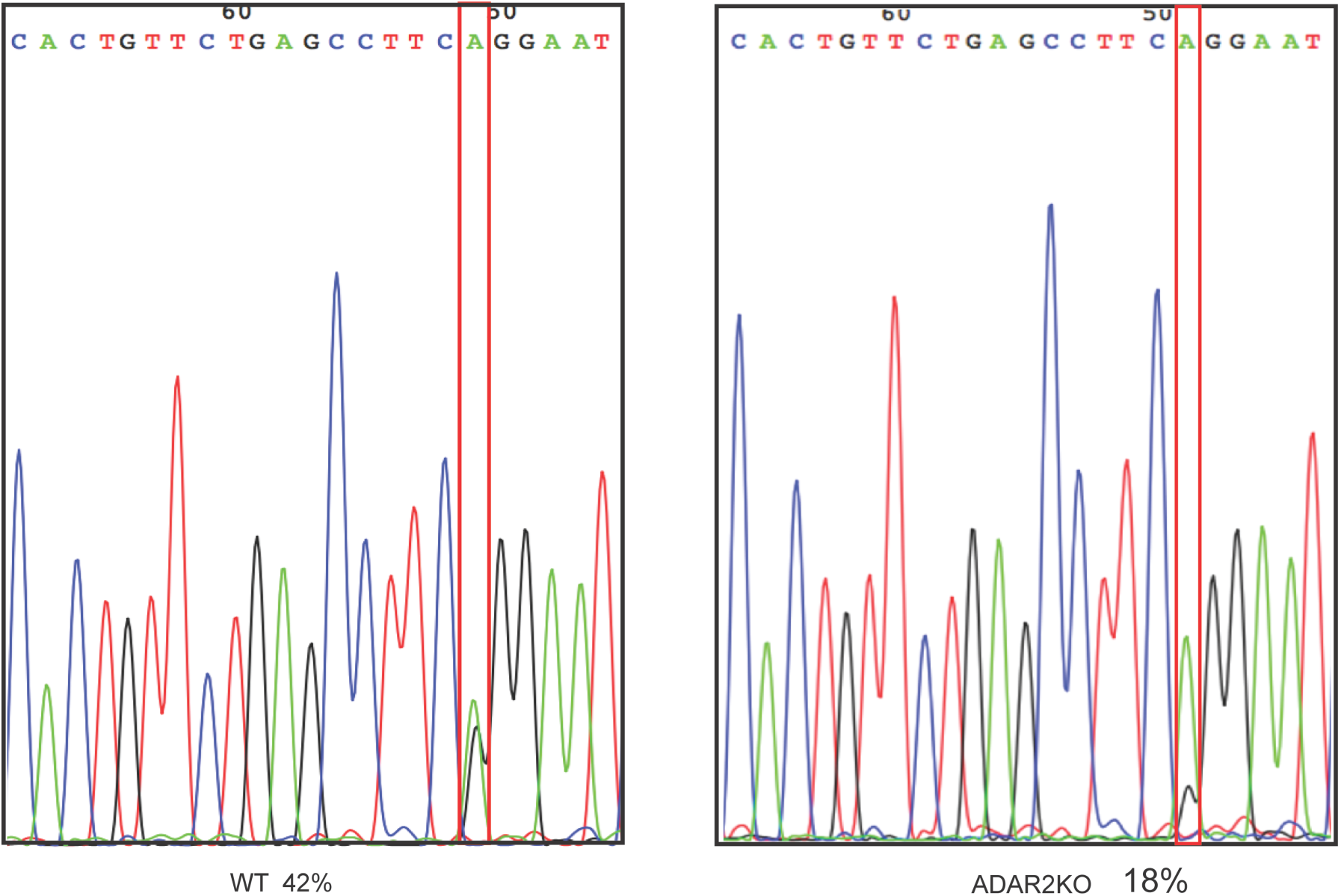
FLNB editing in ADAR2 KO mouse: Electropherogram showing 24% decrease in editing as compared to WT mouse

To our surprise, we did not observe any change in expression of FLNB in the ADAR2^-/-^ mice heart as compared to wild type mice heart (Figure 5). This indicates that edited FLNB might have a specific function.

**Figure 5.**
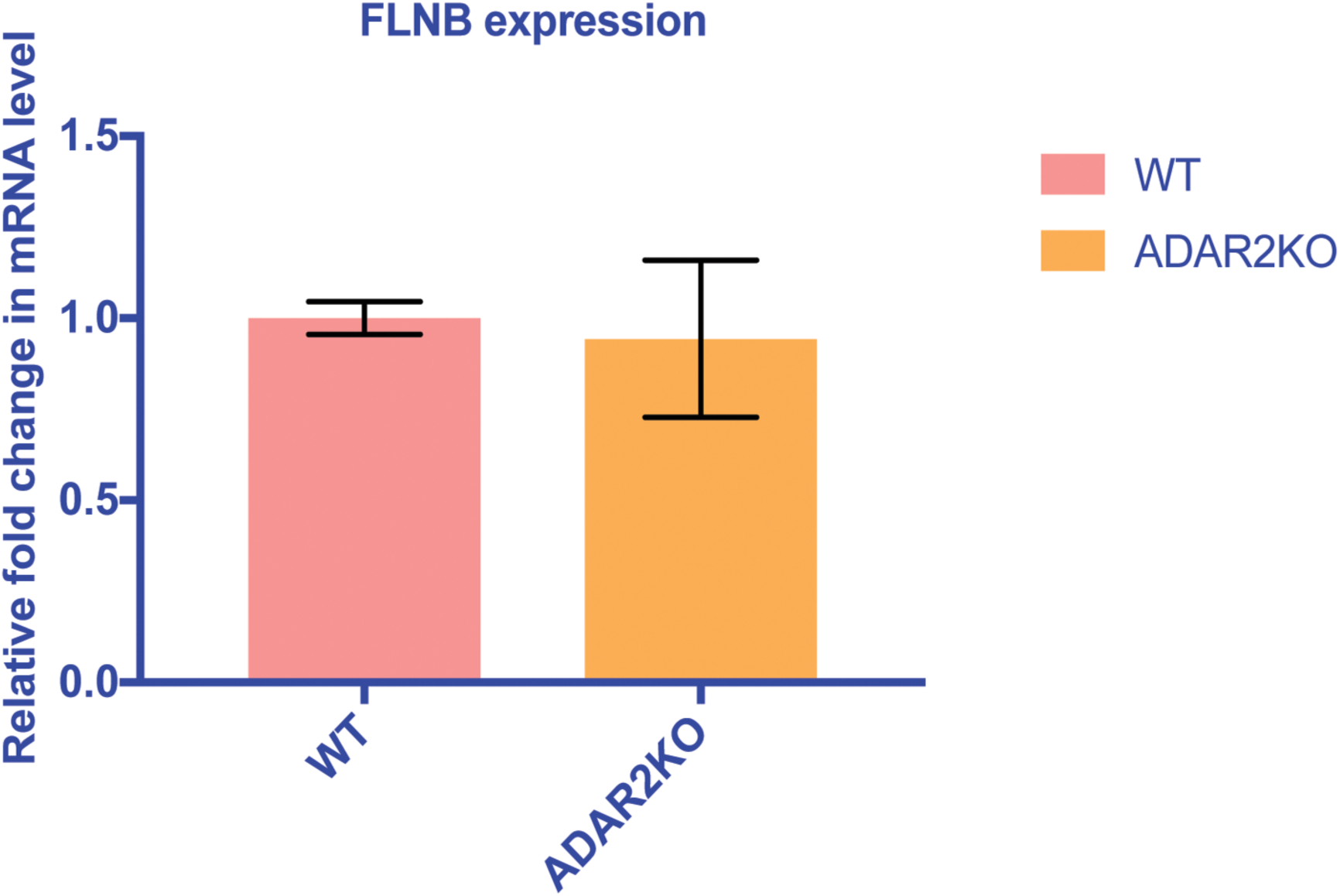
FLNB expression in the ADAR2 KO mouse heart: Bar graph showing no significant change in expression of FLNB in the heart on ADAR2

### ADAR1 expression in ADAR2 KO mouse heart

Since increase in ADAR1 has been found in CHD patients. We determined whether this observed increase in ADAR1 is due to deregulation of ADAR2. Therefore, we determined ADAR1 level in the absence of ADAR2. We did not find any significant change in ADAR1 level (Figure 6). Therefore, we can conclude that the observed increase in ADAR1 is soley because of the CHD defect.

**Figure 6.**
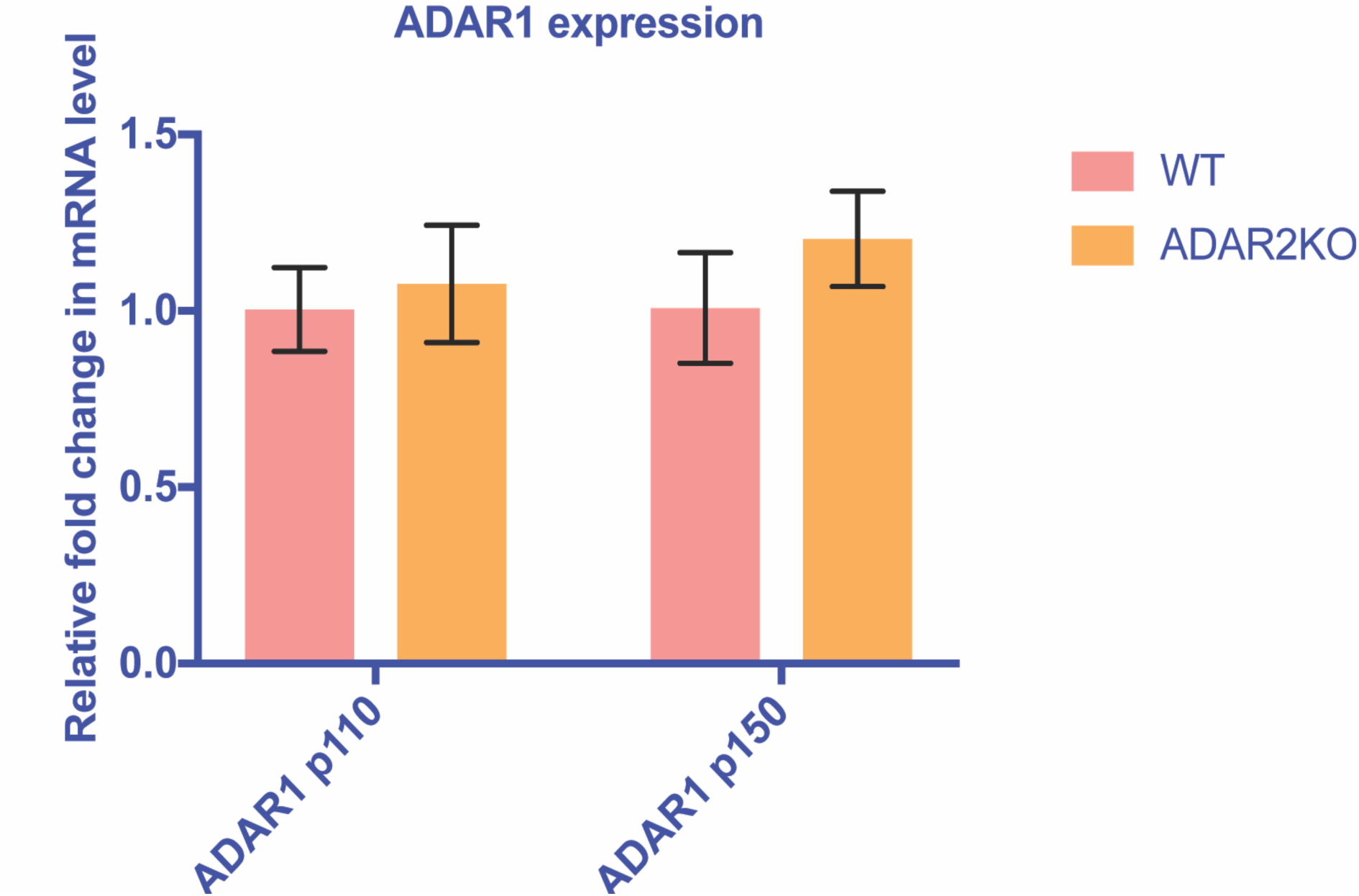
ADAR1 isoforms expression in ADAR2 KO mouse: Bar graph showing no significant change in expression of ADAR1 isoforms (p110 and p150) in the heart on ADAR2

### ADAR2^-/-^ heart shows down-regulation of miR-29b

A previous study focusing on ADAR2 ^-/-^ mice showed a statistically significant decrease in heart rate (15). We performed RNA sequencing of ADAR2^-/-^ mice heart samples in triplicates and observed approximately ~2-fold decrease in miR-29b level consistently at P6 stage. Approximately 1.5 fold down-regulation has been observed for miR451-b, miR451-a, miR19b, (Table 1). To our surprise, we did not observe any up-regulated miRNAs. Quantitative trait loci (QTLs) associated with miR-29 a and b show their potential involvement in cardiac diseases (29). miR-29 family shows strong expression in lung, kidney and the heart. It expresses predominantly approximately 5-12 folds in cardiac fibroblasts as compared to cardiomyocytes. Moreover, the miR-29 family is down-regulated in fibrotic scars after myocardial infarction and can lead to cardiac fibrosis by boosting collagen expression. miR29-b has an antifibrosis role as it targets promoters of several extracellular matrix genes (30). Recent reports have documented a cardioprotective role of miR29-b. miR 29b inhibits angiotensin II induced cardiac fibrosis by targeting TGF- ß/Smad3 pathway (31).

After miR-29b, miR-451(a and b), miR-19b1and different members of let-7, family also showed down-regulation in ADAR2 knockout mice heart However, they showed significant but small down-regulation of only about 1-1.5 fold (Table −1). Aberrant expression of let-7 family has been linked to diverse cardiovascular diseases such as fibrosis, hypertrophy, dilated cardiomyopathy (DM), myocardial infarction (MI), atherosclerosis and hypertension (32). This down regulation of microRNAs on ADAR2 knock-out point towards a potential regulatory mechanism mediated by ADAR2 in the heart development and physiology.

**Table 1.**
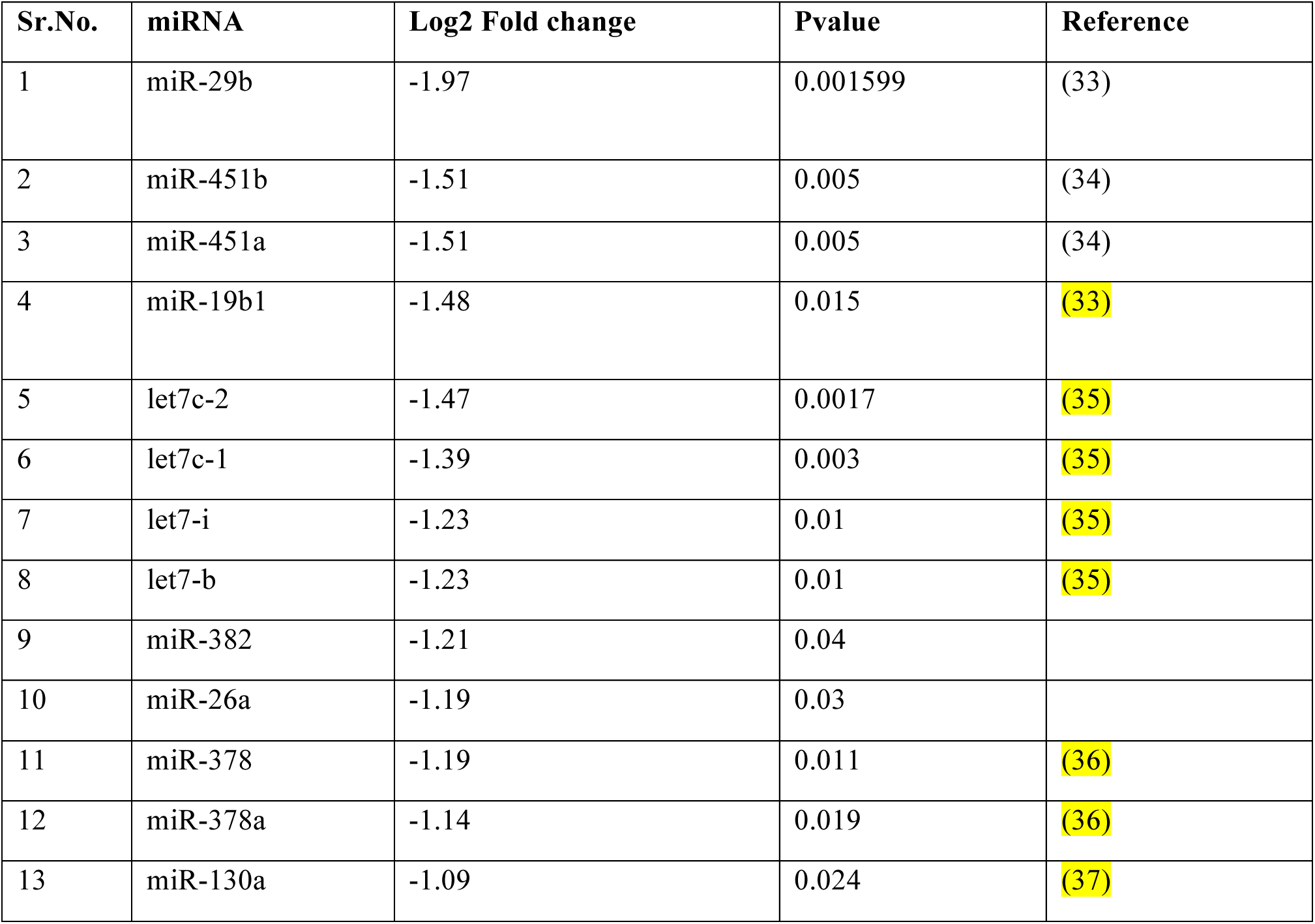
miRNAs significantly downregulated in ADAR2^-/-^ mice heart as compared to wild type

## Discussion

Adenosine deamination by ADARs is a post-transcriptional event that can diversify the transcripts both at sequence as well as structure level. The deregulation of editing has been associated with number of diseases (8, 13, 33).

ADARs play a significant role in development. Moreover the tissue and site specific editing largely affects the differential expression of substrate transcripts (38, 39). In our study, we found a strong down-regulation of ADAR2 and a strong increase in ADAR1 in ASD and VSD patient samples. This observation is in line with previous finding demonstrating increased CTSS mRNA editing due to up-regulation of ADAR1 in human atherosclerotic plaques (14). Since the expression analysis was performed only on the PBMCs from normal and CHD patients we further extended our study to myocardial tissues. Expression analysis of myocardial tissues from ischemic and non-ischemic patients showed a significant decline in ADAR2 expression level (Figure 3). This result supported our finding that ADAR2 not only down-regulates in PBMCS but also showed decreased expression in human cardiomyopathy tissues.

The stong downregulation of ADAR2 with respect to heart disease urged us to further explore what heart related processes might be ADAR2 regulating? To address this query, we used ADAR2 knock out mouse. We chose FLNB which plays an essential role in the heart however lack of ADAR2 strongly decreased its editing. We did not observe any change in FLNB expression in the absence of ADAR2. The increase on ADAR1 in the PBMCs and the simultaneous downregulation of ADAR2 posed a question whether the observed effect is because of ADAR2 or is it the consequence of CHD? We observed no significant change in expression of the ADAR1 isoforms in the absence of ADAR2. This shows that CHD might be triggering an inflammatory response leading to increase in ADAR1. The elevated ADAR1 expression in atherosclerosis has been documented previously (14).

ADARs can modulate microRNA processing and also are capable of retargeting the microRNA to different substrate (35). Like ADAR1, ADAR2 also can modulate microRNA processing. Since a number of micro RNAs like miR-1, miR-423 are associated with heart disease we thought of investigating the microRNA profile in ADAR2 knock out mouse heart (40). Surprisingly, we did not observe any up-regulated micro RNAs in the ADAR2^-/-^ mice heart. However, we observed a decline of ~1.5-2 fold in miR-29b, miR-451, miR19 and members of let −7 family (Table 1).

miR-29b family regulates a plethora of proteins at RNA level that are involved in cardiac fibrosis. This family has highest expression in the heart fibroblast population and comprises of three members miR-29a, miR-29b and miR-29c. miR-29b differs only by one base from miR-29a and miR-29c. Among the three members, miR-29b expresses strongly in cardiac fibroblasts as opposed to miR-29a and miR-29c.(30). Angiotensin-II (Ang-II) induced hypertensive cardiac fibrosis in-vivo and in-vitro is associated with cardiac fibrosis. miR-29b inhibits activation ERK1/2 by preventing its phosphorylation. miR-29b suppresses the TGF-β /Smad-ERK-MAP kinase crosstalk (31). Two members of miR-451 a and b show down-regulation by 1.5 fold. miR-451is down-regulated in the heart tissues from hypertrophic cardiomyopathy patients. It directly targets tuberous sclerosis complex 1 (TSC1) and inhibits formation of auto-phagosomes and cardiac hypertrophy (41). Another microRNA, miR-19b belongs to miR-17-92 cluster. The decreased expression of miR-19b is correlated with cardiovascular diseases (42). miR-19b overexpression promotes differentiation by stimulating cell proliferation and inhibiting Wnt-β-catenin pathway consequently blocking apoptosis of cardiac P19 cells(43).

Apart from the above mentioned microRNAs, some members of let7 family like let7-c, let-7-i and let7-b are down-regulated approximately 1.2-1.4 fold. The human let-7 family comprises of 13 members. Let7-c is elevated in endothelial to mesenchymal transition (EndMT) (32). It shows reduced expression in coronary artery disease patients as opposed to control (44). Let7-i limits the toll like receptors like TLR4 by targeting it. (45) and is down-regulated in dilated cardiomyopathy (32).

Most of the down-regulated microRNAs on ADAR2 knock down in our study are related with cardiovascular disorders. This implies that ADAR2 might have a cardio-protective function as these microRNAs are mostly reduced in different cardiac defects. The increase in ADAR1 in CHD cases is in line with previous finding of elevated ADAR1 expression in endothelial cells stress response and atherosclerosis (14).

## Acknowledgments

We thank Peter Seeburg (MPI, Heidelberg) for the kind gift of Adar2^-/-^ knockout mice and Dr.Michael Jantsch (Medical University, Vienna) for supporting the microRNA work. This work was supported by the Higher Education Commission startup grant IPFP/HRD/HEC/2014/1622. We thank Kate Middleton, for reviewing the manuscript.

